# Enhanced antibody-antigen structure prediction from molecular docking using AlphaFold2

**DOI:** 10.1101/2022.12.26.521961

**Authors:** Francis Gaudreault, Christopher R. Corbeil, Traian Sulea

**Affiliations:** Human Health Therapeutics Research Centre, National Research Council Canada, 6100 Royalmount Avenue, Montreal, Quebec, Canada, H4P 2R2

## Abstract

Predicting the structure of antibody-antigen complexes has tremendous value in biomedical research but unfortunately suffers from a poor performance in real-life applications. AlphaFold2 (AF2) has provided renewed hope for improvements in the field of protein-protein docking but has shown limited success for the medically relevant class of antibody-antigen complexes due to the lack of co-evolutionary constraints. Some research groups have demonstrated the usefulness of the AF2 confidence metrics for assessing the plausibility of protein folding models. In this study, we used physics-based protein docking methods for building decoy sets consisting of low-energy docking solutions that were either geometrically close to the native structure (positives) or not (negatives). The docking models were then fed into AF2 to assess their confidence with a novel composite score based on the pLDDT and pTMscore metrics. We show benefits of the AF2 composite score for rescoring docking poses in two scenarios: (1) a more trivial experiment based on the bound conformations of the antibody and antigen backbone structures, and (2) a more realistic test employing the unbound backbone conformations of the binding partners. Docking success rates improved after AF2 rescoring with particular emphasis on early enrichment of positives at the very top of the re-ranked list of decoys. The AF2 rescoring markedly improved classification of positives and negatives in most systems. Docking models of at least medium quality present in the decoy set, but not necessarily highly ranked by docking methods, benefitted most from AF2 rescoring by experiencing large advances towards the top of the reranked list of models. These improvements, obtained without any calibration or novel methodologies, led to a notable level of performance in antibody-antigen unbound docking that was never achieved previously.

## Introduction

Three-dimensional structures of antibody-antigen complexes can be predicted computationally using physics-based protein-protein docking methods (1–5). Recent benchmark studies have shown that reconstituting the complex by docking the antibody and antigen structures separated from the complex, while keeping their protein backbone conformations as in the bound state, is now a relatively trivial task and mostly a solved problem (4,6). However, antibody-antigen docking remains significantly challenging in the more realistic scenario in which the backbone conformations of the antibody and antigen structures deviate from their bound-state conformations. In this so-called “unbound docking”, the best-ranked model achieves no more than 20% success in predicting a complex structure reasonably close to the native structure (4,6). Encouragingly, these docking methods are relatively good at sampling various binding modes and able to enrich with native-like docking solutions a relatively small fraction of the best-scored 100-1,000 poses out of billions of theoretical ones. The general failure of protein-protein unbound docking thus appears to be caused not only by the high dimensionality of the search space associated with backbone sampling (7), but also to an inability to accurately score and rank docked structures that deviate substantially from the native geometry of the complex. The problem is further exacerbated in modeling antibody-antigen complexes due to the antibody CDR-H3 hypervariable loop which is capable of exploring various backbone conformations in the unbound versus bound states (8,9). In our view, a step forward in the field of protein-protein and antibody-antigen docking would be to employ a complementary method to improve the scoring component. Hence, an objective of this study was to explore a way to “rescue” native-like docking structures and top-rank them ideally among the 1-5 best scoring solutions.

With the release of AlphaFold2 (AF2), artificial intelligence and deep-learning have made a breakthrough towards addressing the protein folding problem (10). Undoubtedly, AF2 has delivered an astonishing performance in the recent blind competition CASP14 by outperforming the previously considered state-of-the-art physics-based protein structure prediction methods (11). One interesting feature of AF2 structure predictions is that they come with confidence levels for different regions of the modeled protein structure. Most of the success and boost in performance of AF2 over its predecessor AlphaFold (12) can be attributed to the inclusion of a Multiple-Sequence-Alignment (MSA) module in addition to the Structure module. The sequence co-evolutionary information that comes with the MSA module embeds structural patterns that strongly influence the thermostability of proteins in general (13) and also of antibodies (14). Thus, providing co-evolutionary information has become a key element towards accurately predicting how proteins fold or arrange themselves within multi-protein complexes (15–17).

Many groups have used AF2 to predict the structure of complexes with the aid of co-evolutionary data. One simple approach consists in tweaking AF2 by adding a long linker between the interacting proteins. Independent releases of AF2 were developed for that specific purpose and were shown to be promising (18,19). For instance, the performance of AF2-Multimer was shown to significantly outperform the traditional physics-based approaches in protein-protein docking (19) when looking only at the top-scoring model. With the aid of AF2, unbound docking has made a huge step forward for those complexes that co-evolved together. However, the requirement for co-evolution data for multimeric units limits its use for predicting antibody-antigen complexes. The strong binding of antibodies towards their antigen is not a result of co-evolution but a result of somatic hypermutation and affinity maturation (20). On this note, a study has shown that the performance of AF2-Multimer against antibody-antigen complexes is no better than the one of physics-based methods (21), indicating the relevancy and need of physics-based methods. While other deep-learning technologies that were developed circumvent the need of the MSA component (22,23), their performance in antibody-antigen structure prediction still remains unproven.

Several examples have already been reported in the scientific literature in which physics-based methods were combined with deep-learning technologies for improved accuracies. On one side, some research groups have used AF as a pre-filter module for generating better input structures for physics-based methods, which showed improved accuracies in folding (24) or docking (25) models. On the other side, others have used it as a structure post-filter module. For instance, one group has used AF2 to assess the plausibility and quality of protein folding models generated by Rosetta (26). An important aspect of their study was bypassing the MSA module. By doing so, no co-evolutionary data was used in the confidence scoring, hence the resulting confidence metrics were based solely on the structure template from the Structure module alone. The authors showed significantly better discrimination of native versus non-native protein structures when folding models were re-ranked using structure-only AF2 confidence metrics relative to the Rosetta physics-based scoring function. These remarkable results obtained with an AF2 version lacking co-evolutionary information opened new possibilities for improving scoring protein structure models generated by physics-based methods.

Our present study, which focuses on the structure prediction of antibody-antigen binding for which no co-evolutionary data exists, is thus inspired and builds upon the aforementioned protein folding study of Roney *et al*. (26). Here, we assessed the plausibility of antibody-antigen interaction models generated using protein-protein docking methods. We investigated if there are benefits in using AF2 in discriminating near-native structures from challenging decoy structures. To ensure consistency, the decoys were generated using three docking methods that are complementary in nature: ProPOSE, a direct-search approach that employs stiff potentials (4), and ZDOCK and PIPER, two Fast Fourier-Transform (FFT)-based approaches employing soft potentials (2,3). Two sets of decoys were generated containing structures with bound and unbound backbone conformations. The “bound-backbone” set of structures, used as control, allowed us to assess the performance of AF2 in a scenario generally more trivial for docking and scoring. The “unbound-backbone” set of structures, on the other hand, is more relevant for real-life applications but considerably more challenging.

## Methods

### Antibody-antigen systems

The antibody-antigen complexes used in the present study are from the union of the protein-protein benchmark version 5.0 dataset (27) and the antibody benchmark dataset (4) originally collected from SAbDab (28). This led to a set of 231 complexes, which we call here the “bound-backbone” set. For 25 antibody-antigen systems from this set we also found crystal structures of the unbound partners, and these structures were collected into the so-called “unbound-backbone” set. The side-chains of the complexes in the bound-backbone set were repacked using SCWRL4 (29). For each of the bound-backbone and unbound-backbone sets, we generated 100 docking models per system with each of the three docking methods described below. Symmetric units were detected by sequence alignments and assembled using PyMOL (30).

### Docking methods

ProPOSE (4) distributed version 1.0.2, ZDOCK (2) version 3.0.2 and PIPER (3) version 0.0.4 were employed to perform the docking simulations. Initially, all input molecules were repaired (addition of missing side-chains), prepared (removal of small molecules, addition of hydrogen atoms, addition of capping groups where needed) and then charged and energy-minimized using the Amber force-field (31) and suite of tools (32). Hydrogen atoms and capping groups were removed prior to performing docking with ZDOCK and PIPER for incompatibility reasons. ClusPro (33) is often used to post-process the top 1,000 predicted models generated by PIPER. Since our objective was to generate a sufficiently high number of energetically-favorable decoy models with each docking method, we dropped ClusPro for the following reasons: a) redundancy with PIPER models, b) small number of generated decoys due to clustering and c) difficulty of attributing a score to each cluster representative model.

Docking was directed to the complementarity-determining region (CDR) of each antibody. The CDR boundaries were defined using the Kabat numbering scheme (34). Restriction to the CDR region was enforced for each of the docking methods as the following. For ProPOSE, the atoms within the CDR region were labeled with the HITSET flag. For ZDOCK, all atoms outside the CDR region were forced to have the atom type 19. For PIPER, a mask of 1 Å radius around all CDR atoms was applied.

### AlphaFold2 preparations

The multiple chains from the antibody-antigen docked models were connected into single chains by using 50-residue-long glycine linkers. The residues were renumbered in sequential order. The merged 3D structure was used as template for AF2 modeling. Hence, AF2 was provided as input the docked 3D template and the concatenated single-chain or “monomer” sequence for extraction of features data. The *template_all_atom_masks* input feature was used to apply an inclusive mask to the backbone and Cβ atoms to indirectly strip all side-chains from the template structure and artificially mutate all amino acids to alanine. Glycine residues were added the missing Cβ atoms. The *template_sequence* and *template_aatype* features are the alanine sequence and one-hot encoded alanine sequence of the template, respectively. The *template_domain_names* feature was defined as none. The multiple sequence alignment data used to build the monomer features were blanked and only contained the full sequence of the merged chains.

The AF2 calculations were run on NVidia A100, P100 Pascal and V100 Volta cards. AlphaFold version 2.2.2 was used across computing clusters. The model model_1_ptm was used throughout the study while all other AF2 parameters were kept to the default values.

### Scoring schemes

From the antibody-antigen models predicted by AF2, confidence metrics were extracted at every position in the sequence. The scores pLDDT and pTMscore average the confidence metrics over the entire antibody-antigen models and report the confidence score as a singular value. The value typically falls within the range 0 (not probable) to 1 (highly probable). These scores were used to derive the AF2 composite score used in ranking and defined according to Equation 1.

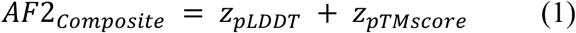

The *z*_pTMscore_ and *z*_pLDDT_ are standardized scores (or Z-scores) that are calculated from an ensemble of docked models for one given system. Higher AF2_Composite_ scores are indicative of higher confidence relative to the ensemble.

The energetic scores outputted by ProPOSE, ZDOCK and PIPER were also collected and used in ranking the models. To be consistent with the values reported by AF2_Composite_, for ProPOSE and PIPER, the sign of the scores were flipped such that higher scores are more favorable energetically. The scores outputted by docking methods cannot be compared in absolute units across docking methods. With the objective to compare and eventually aggregate docking poses and scores obtained from various docking methods, a standardization scheme was also applied to the energetic scores calculated in the ProPOSE, ZDOCK and PIPER docking runs. While the standardization is flawed as it assumes docking methods produce similar distribution of scores, we believe it is a good compromise and is how one would go about merging docking methods.

## Results

### Decoys sets

Antibody-antigen docking simulations were performed with the ProPOSE (4), ZDOCK (2) and PIPER (3) methods using as input the protein structures in their bound-backbone and unbound-backbone conformations (see Methods). The top 100 models predicted by each method were collected. In total, 68,671 and 7,440 models were generated from 231 and 25 unique systems for the bound-backbone and unbound-backbone decoy sets, respectively (**Table 1**).

**Table 1.** Composition of the bound-backbone set and unbound-backbone set of antibody-antigen complexes used in this study.

| Decoys set | Method | Systems <sup>a</sup> | Models <sup>b</sup> | Positives (%) <sup>c</sup> | Negatives | P/S <sup>d</sup> |
| --- | --- | --- | --- | --- | --- | --- |
| Bound-backbone | ProPOSE | 231 | 22515 | 974 (4.3%) | 21541 | 4.2 ± 4.3 |
|  | ZDOCK | 231 | 23056 | 740 (3.2%) | 22316 | 3.2 ± 5.9 |
|  | PIPER | 231 | 23100 | 2996 (13.0%) | 20140 | 13.0 ± 21.9 |
| Unbound-backbone | ProPOSE | 25 | 2500 | 34 (1.4%) | 2466 | 1.4 ± 2.2 |
|  | ZDOCK | 25 | 2500 | 63 (2.5%) | 2437 | 2.5 ± 3.9 |
|  | PIPER | 25 | 2440 | 236 (9.7%) | 2204 | 9.4 ± 17.0 |

<sup>a</sup> Systems for which the docking method produced results without technical error.
<sup>b</sup> Docking methods do not necessarily output 100 models even when requested by the user.
<sup>c</sup> Based on the AlphaFold2-reconstructed template and models of at least acceptable quality according to CAPRI classification (35).
<sup>d</sup> Mean and standard deviation for the number in positives per system (P/S).

Models were compared to their respective crystal structures and attributed a quality as previously described in the CAPRI blind challenges (35). The models with high, medium or acceptable quality were reported in the “positives” set as opposed to the incorrectly predicted models reported in the “negatives” set. The various docking methods produced variable fractions of positives (1−13%) for a total of 4,710 and 333 positives, which came along with 63,997 and 7,087 negatives, for the bound-backbone and unbound-backbone sets, respectively. On average, 2−13 positives per system were generated. The uncharacteristically high numbers for positives per system generated by PIPER are due to redundancy within the top-100 models, which are typically subsequently clustered with the ClusPro method (33).

### Quality of AlphaFold2-generated models

AlphaFold2 was run providing as input the amino-acid sequence and the structural template of each of the docking-generated model in the two sets. Noteworthy, only the backbone structure of the template was provided, *i*.*e*. the template was stripped of all its side-chains similarly with a previous study on protein folding (26). This procedure allows the release of unnecessary constraints to the Structure module and forces a rebuilding of the side-chains by AF2. We compared the AF2-generated model to its provided template to assess the amount of structural changes at the antibody-antigen interface that occur from rebuilding. Specifically, the fraction of conserved contacts and interface RMSD were calculated and used as proxy for assessing structural divergence (**Figures 1A** and **1B**). On average, 52% of the template contacts were retained upon remodeling. In terms of interface RMSD, the structures deviated by 1.25 Å on average. These values were maintained in the unbound-backbone set (**Figure S1**). These structural discrepancies underline the inability of AF2 to accurately reproduce the fine atomic details at the interface from its lack of physics, as shown previously (10).

**Figure 1.**
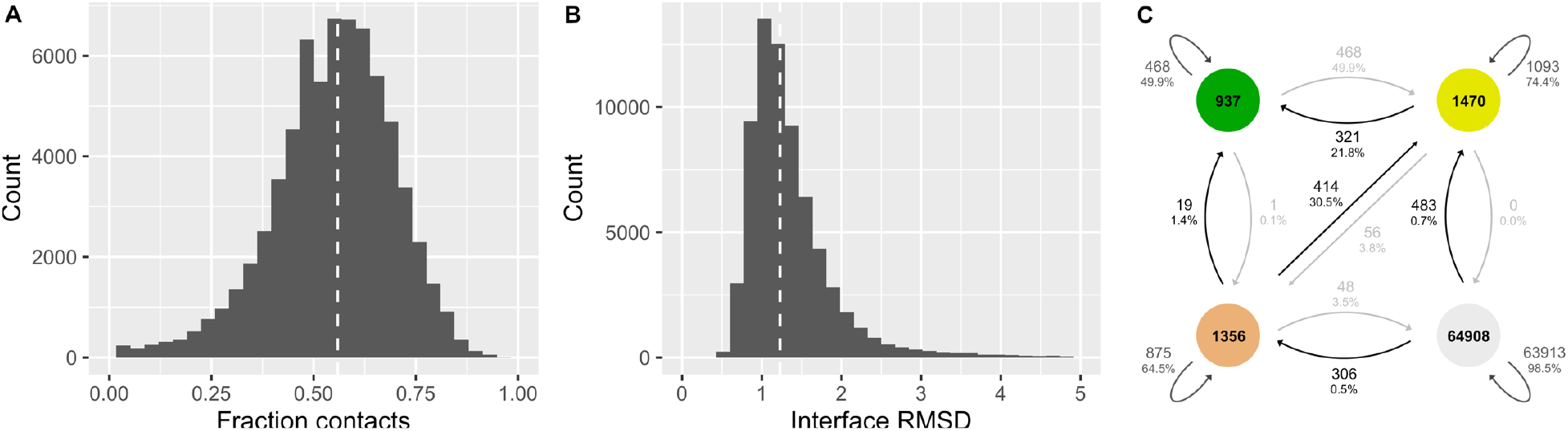
Quality assessment of the AlphaFold2-generated models in relation to its provided docking-generated template structure. Distribution in the (**A**) fraction of conserved template contacts and (**B**) interface RMSD between the AF2-generated model and its provided docking-generated template for the decoys in the bound-backbone set. The RMSD calculations only include the Cα atoms. All decoys generated with ProPOSE, ZDOCK and PIPER were combined. The median of the distribution is indicated by the dashed white line. (**C**) Transitions from the docking-generated models to the corresponding AF2-generated models in terms of model quality relative to the crystal structure. Transitions between the high-quality and incorrect classes were removed for clarity. The transition from incorrect to high quality occurs in 206 instances (0.3%). Colors denote structure quality levels as defined by CAPRI classification: high (green), medium (yellow), acceptable (beige) and incorrect (grey). Figures in this paper were produced in R (40) using the ggplot2 library (41).

From the fraction of template contacts alone, it remains unclear if such changes in interface contacts result in a loss or a gain of contacts relative to the ones observed in the experimentally-resolved structure. Therefore, we then compared the model quality of the docking-generated and AF2-generated structures to the experimental structures for the bound-backbone (**Figure 1C**) and unbound-backbone (**Figure S1C**) sets. In general, most models produced by AF2 retained the quality of their docking-generated templates. In absolute terms, there are more positives in AF2-generated models than docking-generated models, with 4,710 and 333 positives as opposed to 3,763 and 271 positives, for the bound-backbone and unbound-backbone sets, respectively. This prompted us to use the AF2-generated models for success evaluation in the sections that follow.

One can note a relation between the ability of AF2 to increase model quality and the quality of the docking template used as input. High-quality docking models tend to lose the most of their quality, in part due to the stringency of the high-quality metrics paired with the inability of AF2 to reproduce physics at atomic level. In net numbers, there were 937 and 1,014 positives of high-quality for docking-generated and AF2-generated models, respectively. Benefits of AF2 in producing better-quality models were seen mainly for docking templates of medium and acceptable qualities, with increases of 22% and 31%, respectively (**Figure 1C**). These trends were maintained in the unbound-backbone set, with increases of 10% and 20%, respectively (**Figure S1C**). It is encouraging to see improvements even for the models of acceptable quality, which are considered to be at the limit of practical applicability.

### AlphaFold2 rescoring improves true-positive ranking

The area-under-the-curve (AUC) of the receiver-operating-characteristic (ROC) curve was calculated per each system to evaluate the overall classification of true positives and false positives. The AF2_Composite_ score was used in re-ranking antibody-antigen docking models. The AUCs of models ranked by docking scores were compared to the AUCs of models ranked by the AF2_Composite_ score for those systems with at least one positive model (**Figure 2**). On average, the AUCs were significantly improved after rescoring with AF2_Composite_, and improvements were obtained for all three individual docking methods.

**Figure 2.**
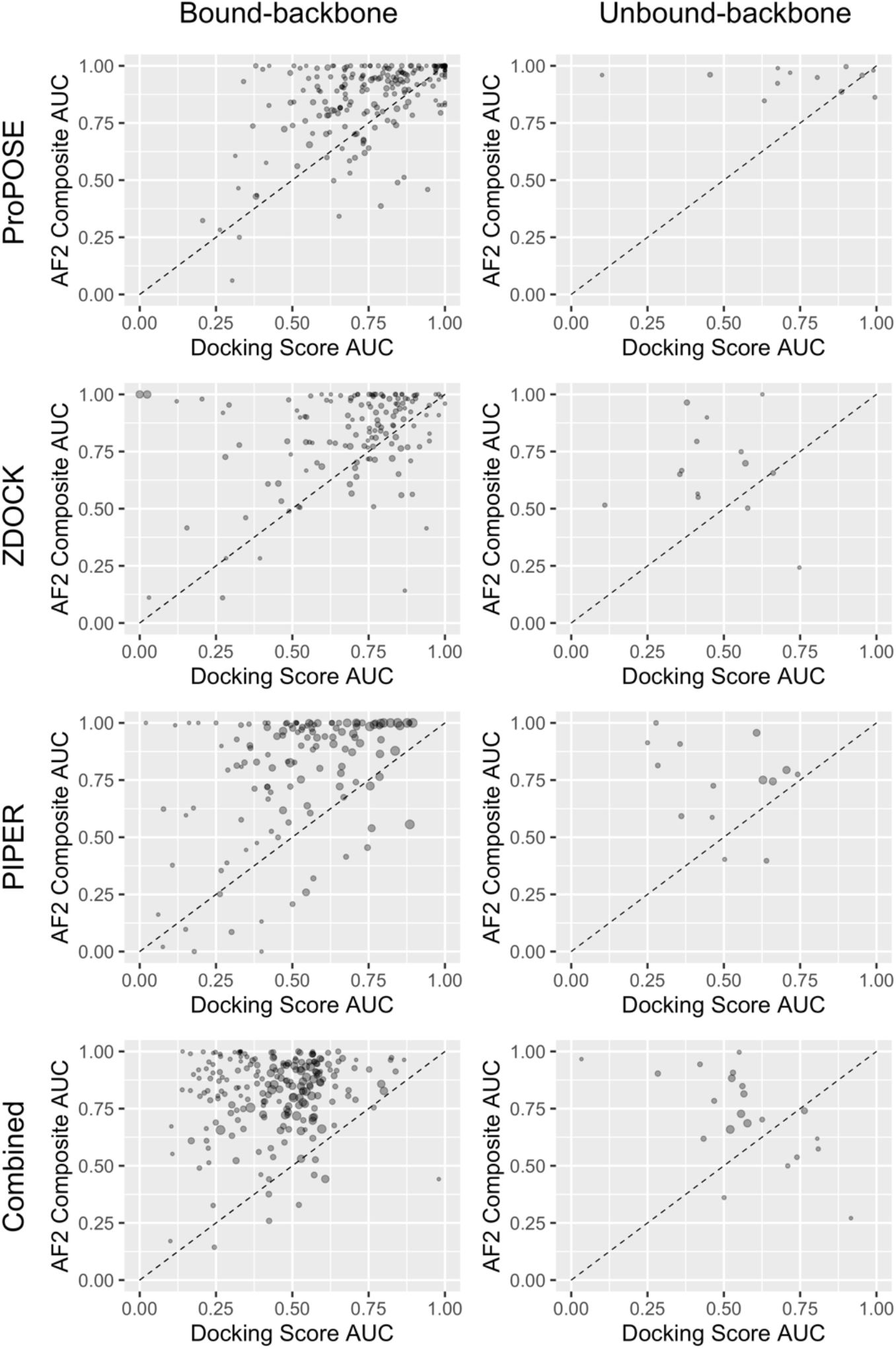
Classification of true positives and false positives according to AUC for the ROC curves based on docking scores relative to AF2 composite scores. Each point in the plot represents a different system. The points are area-weighted by the number of corresponding positives on a 1-to-10 scale. The diagonal line is drawn to indicate the impact of AF2-rescoring. The number of data points (antibody-antigen systems) in each plot are 208, 12, 137, 14, 127, 14, 218 and 21 (from left to right and top to bottom). AF2 rescoring improves the classification for 145 (70%), 10 (83%), 104 (76%), 11 (79%), 111 (87%), 14 (86%), 209 (96%) and 14 (67%) antibody-antigen systems. Positives include high, medium and acceptable quality models according to CAPRI classification (35).

In the bound-backbone set, AF2_Composite_ improves the classification for 145 (70%), 104 (76%) and 111 (87%) systems having at least one positive model for ProPOSE, ZDOCK and PIPER, respectively. ProPOSE benefits the least from AF2 rescoring given its high accuracy in bound docking, hence an average AUC improvement of only 0.10 was obtained. Other docking methods benefitted more from the AF2 treatment, with average AUC improvements of 0.13 and 0.27 for ZDOCK and PIPER, respectively. When the acceptable-quality poses were removed from the definition of success (**Figure S2**), AUC improvements increased for both ZDOCK (0.19) and PIPER (0.32) as noted by the higher density of points in the upper section of the plots, suggesting that poorer quality models were harder to score with high confidence by AF2. Still, it is impressive to see systems with poor or random AUC (< 0.6) based on docking scores improve to remarkable levels (AUC > 0.9) after AF2_Composite_ rescoring; these included 12 systems (9%) for ZDOCK and 35 systems (28%) for PIPER.

In the unbound-backbone set, the classification is improved for 10 (83%), 11 (79%) and 14 (86%) systems having at least one positive model for ProPOSE, ZDOCK and PIPER, respectively (**Figure 2**). The average AUC improvements in this set were larger than in the bound-backbone set, reaching 0.21, 0.20 and 0.24 for ProPOSE, ZDOCK and PIPER, respectively. When acceptable-quality models were removed, average AUC improvements were 0.14, 0.33 and 0.38, respectively.

We also analyzed the combined set of decoys from all three docking methods (ProPOSE + ZDOCK + PIPER) to assess the ranking ability of AF2_Composite_ when challenged with models obtained from different physics-based methods which may slightly vary in their protein preparation. The AUCs were markedly improved after AF2 rescoring for 209 (96%) and 14 (67%) systems for the bound-backbone set and unbound-backbone set, respectively. In 70 out of 218 bound-backbone systems (32%) and 5 out of 21 (24%) unbound-backbone systems with poor docking score-based AUCs, remarkably high AUC values were obtained after AF2 rescoring.

### AlphaFold2 rescoring improves early success rates

The AUC is a global indicator of performance and is often insufficient to capture important early success in terms of true positives enrichment. Having better AUC from rescoring should normally translate into higher success rate in the top fractions, but it remains unclear how many poses have to be inspected by the user to get one successful model. For this reason, success rates were calculated along the ranked list of models and plotted on a logarithmic scale (**Figure 3**). We distinguish late success rates (top 100) from early rates (top 1 and top 5). Here, the top-5 success rates were used as it is generally a tractable number of models to inspect. The success rates based on docking scores were again compared to those obtained after re-ranking with the AF2_Composite_ score.

**Figure 3.**
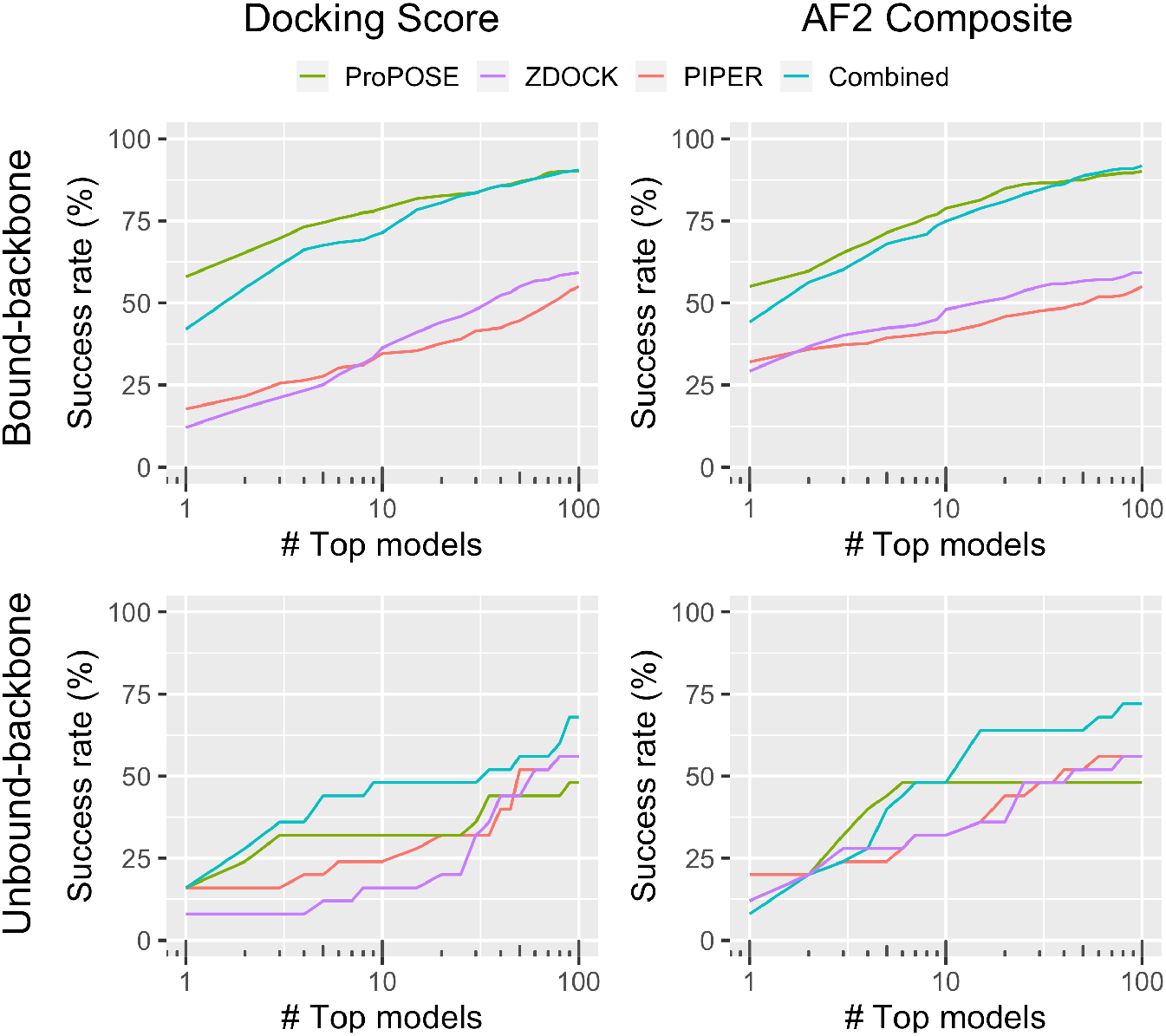
Success rates for various docking methods followed by AF2 rescoring as a function of the number of top-ranked models. The rates are shown for models ranked using the docking scores and after rescoring with the AF2_Composite_ score for the bound-backbone set and unbound-backbone set. The success was evaluated using the AF2-reconstructed models. An acceptable-quality model according to CAPRI classification (35) was the minimum requirement for success.

In the bound-backbone set, the success rates plateaued near the top-50 mark at 90%, 59% and 55% levels for ProPOSE, ZDOCK and PIPER, respectively (**Figure 3**). The success rates were broken down for each model quality in the Supplementary Information (**Figures S3** and **S4**) as well as for success calculated from the docking scores and models (**Figures S5** and **S6**). Rescoring with AF2 helped reaching the plateau earlier, *e*.*g*. for ZDOCK and PIPER the plateau is nearly reached faster. AF2 rescoring also helped with the early success of recovering true positives. While the early success for ProPOSE remained unchanged after AF2 rescoring, the rates at the top-1 model level for ZDOCK and PIPER were markedly improved to reach 29% and 32% from 12% and 18%, respectively. At the level of top-5 models, success rates for ZDOCK and PIPER reached respectively 42% and 39% after AF2 rescoring from 25% and 28% based on docking scores. Overall, it is reassuring to see that AF2 enhanced antibody-antigen docking predictability in this more trivial but still important control experiment, i.e., bound-backbone docking.

In the unbound-backbone set, the trends were nearly identical with those seen in the bound-docking experiment (**Figure 3**). While success rates with the top-1 model remained unchanged after AF2 rescoring, the top-5 models afforded superior success rates superior for all docking methods employed. Hence, success rates of 32%, 12% and 20% were obtained in the top-5 models for ProPOSE, ZDOCK and PIPER, respectively, when ranked with their internal scoring functions. Upon rescoring with the AF2_Composite_ score, these rates went up to 44%, 28% and 24%, respectively. Another way to see this improvement is that a late success rate of 48% that was reached at the top-90 models level with the ProPOSE scoring function was achieved much earlier, at top-6 models only, after AF2 rescoring.

Pulling together of models from all three docking methods followed by AF2 rescoring achieves a success rate of 40% and 48% in the top-5 and top-10 models for unbound docking. These success rates are higher than what was ever achieved in antibody-antigen unbound docking thus far. Further analysis showed that medium-quality models were preferentially enriched over acceptable-quality models upon rescoring with AF2, reinforcing the notion that poorer quality models are harder to score with high confidence by AF2 (**Figure S4**).

### AlphaFold2 composite score can separate positive from negatives

The density distribution of scores for the positives and negatives obtained with the docking methods covered in this study and with the AF2_Composite_ function were comparatively analyzed. For the bound-backbone set, the separation between the means of the distributions of positives and negatives were 1.3, 0.6 and 0.2 standard deviations based on the normalized docking scores from ProPOSE, ZDOCK and PIPER, respectively (**Figure 4**). Thus, the ProPOSE score appears to already discriminate to some extent the true positives from decoy models in the bound-docking scenario. Rescoring with the AF2_Composite_ score increased the separation between positives and negatives for all three methods to 2.5, 2.2 and 1.4 standard deviations. This highlights a stronger overall power in discriminating true positives with the AF2_Composite_ scoring scheme that is less subjective to small perturbations in the structure. The differences in separations achieved with the AF2-reranked models between the various docking methods (PIPER < ZDOCK < ProPOSE) also highlight that docked decoy sets should not be treated equally and suggest an influence of model quality on the achieved AF2 confidence levels.

**Figure 4.**
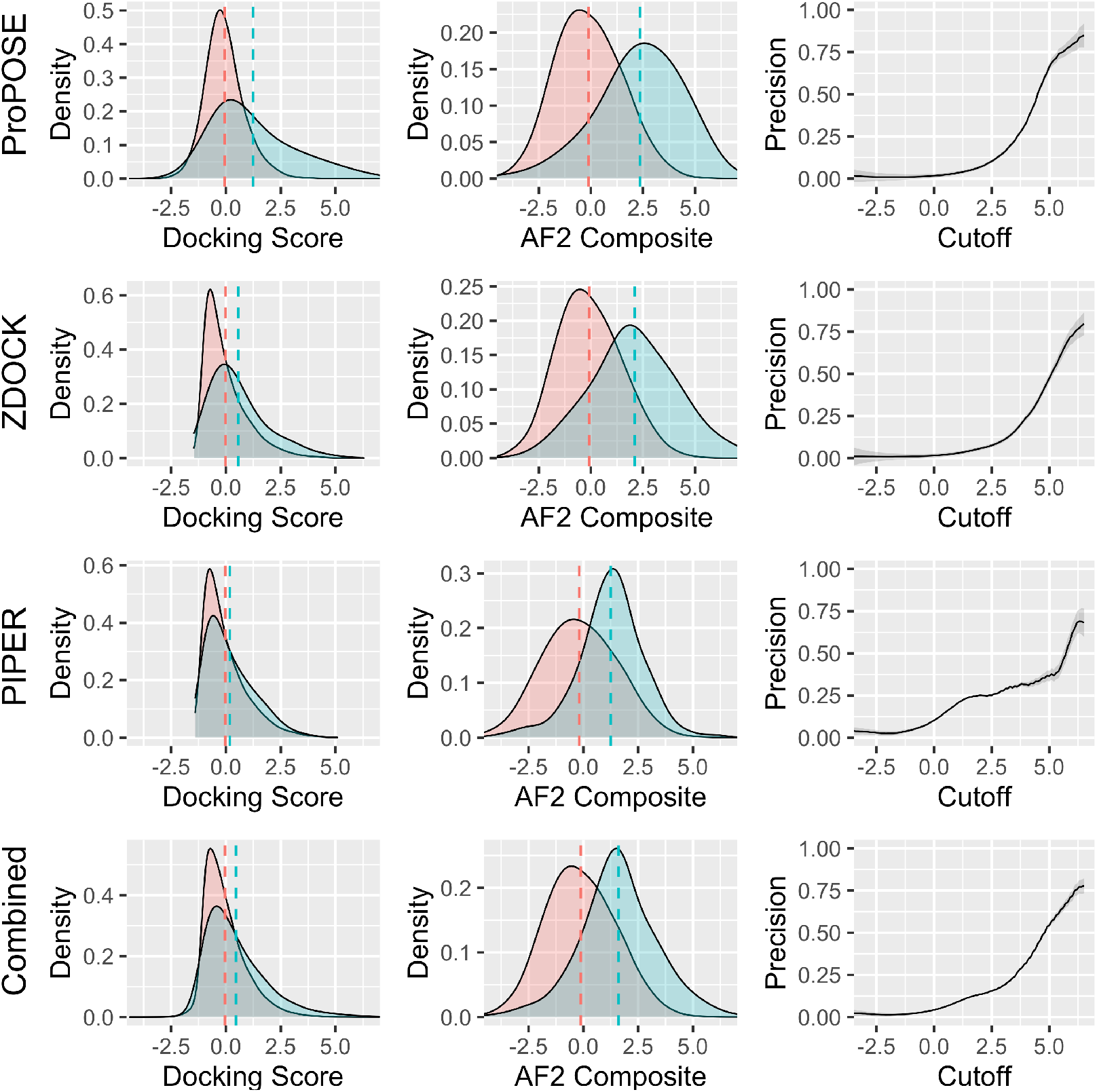
Density plots for the docking scores and the AF2_Composite_ score for the negative and positive sets from the bound-backbone set. The means of the distributions are marked with dashed lines. Precision curves were built from calculating the fraction in the number of true positive over the total number of true positives and false positives according to a given cutoff in the AF2_Composite_ score. The smoothed densities were used to build the precision curves to avoid outlier bias. Shadings around the precision curves indicate the errors on these curves, which were estimated based on the number of data points, *i*.*e*. less data points incur larger errors. For all docking methods evaluated, the likelihood of a true positive event increases with higher cutoff values in the AF2_Composite_ score.

One important property of a metric is its ability to set a threshold from which one can draw confidence levels of success and obtain more valid conclusions. This has implications in knowing, for instance, if a prospective docking run succeeded in generating top-ranked near-native poses or not. To this end, precision curves were built from the smoothed probabilities for positives versus negatives as a function of a sliding threshold for the AF2_Composite_ score (**Figure 4**). These precision curves describe the ratio of true positives found over the number of models sampled at a given threshold of the score. The precision curves are consistent across the decoy sets generated with the different docking methods. For example, a cutoff set at a value of 5.0 for the AF2_Composite_ score suggests a 70% chance of successfully generating a native-like antibody-antigen model when using ProPOSE for docking.

Similar trends were obtained for densities and precision curves with the unbound-backbone set (**Figure S7**). The strong benefit of AF2 rescoring is highlighted by improvements from 1.4, -0.1 and 0.2 to 2.9, 1.3 and 1.2 standard deviations for ProPOSE, ZDOCK and PIPER, respectively, in separating positives from negatives. However, the precision curves are plagued by the low counts of systems in this set. The calculated errors are much higher provided the weaker statistics. If more systems and better-quality structures closer to the native complex could be fed into AF2, most likely better precision curves that approach those seen in the bound-backbone set could be obtained in unbound docking as well.

## Discussion

### AlphaFold2 helps alleviate the scoring problem

In this study, we hypothesized that AlphaFold2 could be used to circumvent the weakness of physics-based docking methods with regards to scoring antibody-antigen complexes. A first assumption was that docking methods, based on their stronger underlying physics, are generally well-suited for finding plausible binding modes for antibody-antigen complexes. A second assumption was that docking scores are generally unable to distinguish true positives from false positives. In short, it was overall assumed that docking methods can be used for the purpose of sampling but not scoring. The AF2_Composite_ score was derived to perform the task of scoring and re-assess the acceptability of the docking-generated models.

The AF2_Composite_ score disregards the predicted energy of the physics-based methods and is solely based on the pLDDT and pTMscore confidence metrics from AF2. These two metrics capture distinct and complementary features for compounding the structural errors. On the one hand, pLDDT is a more relevant metric for local error approximation. On the other hand, pTMscore is a global metric better suited for larger systems such as in the prediction of complexes (10). The normalized values for the pLDDT and pTMscore scores were summed to derive the AF2_Composite_ score. We found this summation to be central for success. In fact, using either metric in isolation led to significant lower performance (data not shown). The AF2_Composite_ detects outstanding models for which the normalized pLDDT and pTMscore metrics agree on high scoring and favorable ranking with high confidence. This agreement is a strong determinant for describing the quality of the model (**Figure S8**). We noted that this agreement weakens as the quality of the docked model deteriorates. For example, pLDDT and pTMscore cross-correlate poorly (R^2^ < 0.4) on models of acceptable quality alone, whereas higher correlations are seen on high and medium-quality models. Our results showing improved classification after rescoring also indirectly point to the same conclusion, that is, that these metrics agree better for positives than negatives (**Figure 2**). The AF2_Composite_ score is a convenient, simple yet effective approach for scoring that is not subjective to any sort of training or calibration.

It is difficult to compare the absolute values for the pLDDT and pTMscore metrics across various protein systems and docking methods, as they are influenced by many factors such as protein preparation, format and input structure. Therefore, the components of the AF2_Composite_ score were standardized, which makes the AF2_Composite_ score system-independent. One drawback of data normalization is the inability to discriminate success; a distribution comprised only of negatives may look identical to one of only positives. However, an important aspect of the AF2_Composite_ is that its distribution is no longer a standardized normal distribution, due to its reliance upon the agreement between its component terms. Incorrectly predicted models have lower agreement in their pLDDT and pTMscore metrics (**Figure S8**) and lower values of those metrics in absolute terms (**Figure S9**). Inversely, medium and high-quality models agree more on their confidence metrics while having higher values in absolute terms. Statistically, an initial distribution composed only of incorrectly predicted poses would suddenly appear skewed as higher quality models are introduced in it. The AF2_Composite_ score can thus be treated in absolute terms and allows the definition of a threshold that can be used to discriminate the docking runs that succeeded from those that failed (**Figure 4**).

### Structure quality of the template matters

Although for some complexes we observed a significant remodeling of the antibody-antigen interface leading to higher-quality models or success, it is not guaranteed that this is a general behavior. In fact, for most complexes, the AF2 remodeled structure retained the quality of the supplied docked model (**Figure 1**). In this study, we used a quite loose definition for success that includes acceptable quality by CAPRI metrics (35), which is standard practice the field. However, model quality has a major influence on the confidence scores. Not only the high and medium-quality models have more value because of higher agreement of normalized pLDDT and pTMscore components, their absolute confidence scores approach those of crystal structures (**Figure S9**). Many systems had binding interfaces identified only broadly (acceptable-quality), which is not always sufficient as those models were predicted with much less confidence.

Given the importance of model quality for high-confidence scores, the docking methods that tend to generate larger proportions of high and medium-quality models should be preferred for rescoring by AF2 instead of those that produce a large fraction of acceptable-quality models. The unbound docking with ProPOSE is one such case which benefitted most from AF2 rescoring and led to the most impressive AUC and success rate results (**Figure 3**). The improvements incurred from AF2 rescoring of ZDOCK and PIPER poses are less striking in that regard. Along these lines, ClusPro processing of PIPER docking poses may also be less suited for AF2 rescoring, since in a previous comparative study ClusPro produced the largest number of acceptable-quality models among all docking methods evaluated (36).

### Influential factors for AlphaFold2 success

The first of several decisions made in this study was the granularity of protein sequence used by AF2 onto the docked template structure, which turned out to be of critical importance. We found that providing a full atomistic template with side-chains added too many constraints to AF2. It prohibited the Structure module of AF2 to work with its own scoring scheme and instead forced to work in the context of another scoring function. Using the full side-chain atomistic model led to significantly worse performance (data not shown). Conversely, forcing the protein sequence of the template to poly-alanine allowed freedom to the AF2 Structure module to remodel the antibody-antigen interface with internally consistent structural determinants for binding, which were necessary for assessing model confidence accurately. Moreover, by allowing AF2 to remodel the binding interface, we simplified the docking task to finding an overall good docking pose among several given backbone geometries. This is an easier problem than having to predict the fine atomic details at the interface, given the size of the search space to be explored.

Secondly, a choice in the final structural models used for AF2 rescoring had to be made. The side-chain reconstructed models produced by AF2 were noted to be different structurally than the templates at fine atomic levels (**Figure 1**). In some cases, these changes were sufficiently large enough to be attributed a different quality, or even different arrangements at the backbone level that could qualify as local “redocking”. One decision that had to be made was which template to use for evaluating success. The overall net sum of positive models was greater for the AF2-reconstructed models when compared to the initial docking-generated ones. Therefore, throughout the study, we hence used the models reconstructed by AlphaFold2. However, a decrease in the number of high-quality structures was noticed for the bound-backbone set due to the less atomistic physical method in AF2. One suggestion for practical use is that the choice of the template should be based on the application. In applications that require clash-free structures with finer atomic details such as affinity maturation, one should employ docking-generated models. For coarser applications like epitope mapping, the AF2-reconstructed models are preferred according to our results here.

Another decision made was related to the number of docking models per system. The AF2_Composite_ score is inevitably influenced by the number of models included per system given that its component terms are normalized (Z-scores). Higher number of models are required to obtain a better approximation for the normal distribution and derive more accurate composite scores (**Figure S10**). For instance, standardized values obtained from a distribution with only 5 models are more likely to have larger errors than on one obtained from 100 models. Including the top 25 poses appears to be sufficient to reduce the error below one unit of AF2_Composite_ score. Therefore, generating more docking models not only leads to an increase likelihood of visiting near-native poses, but also to more accurate confidence scores. Improvement in success would most likely be observed if the top-1000 docking models could be included. However, large number of models per system come with an increased computational cost. If one cannot afford hundreds of AF2 calculations per system, resampling techniques such as bootstrapping (37) should be considered.

One common practice in the field consists in combining many methods for consensus prediction in order to circumvent the limitations and inherent biases from training each method (38,39). Docking methods are no different on that regard, with some dockers being more successful on specific protein systems. By combining multiple methods like ProPOSE, ZDOCK and PIPER, we hoped to also benefit from their orthogonality and increase success rates. While there were significant improvements in global terms (e.g., AUC-ROC), early success rates after AF2 rescoring improved only marginally when all docking methods were combined relative to individual docking methods. Lastly, each physics-based scoring function has specific biases, approximations and constraints in their underlying force-fields, which lead to varying sensitivities to fine structural details. Scoring effectively and fairly models produced from a variety of docking methods is a difficult task in consensus modelling. By using poly-alanine docking templates, we showed in this study that AF2 can be used as an effective platform for rescoring complexes in an unconstrained environment irrespective of the physics-based docking method and without a need for prior training. This is a promising finding for future method developments.

In conclusion, the method proposed in this study appears to hold promise for addressing one of the last largely unsolved major problems in protein modeling, namely, the antibody-antigen docking.

## Acknowledgments

We thank Digital Research Alliance of Canada (formerly Compute Canada) for computing resource allocation for project number 4191. We also thank Dr. Enrico O. Purisima for useful discussions and feedback on this study.

