## Supplementary Information for "Enhanced antibody-antigen structure prediction from molecular docking using AlphaFold2"

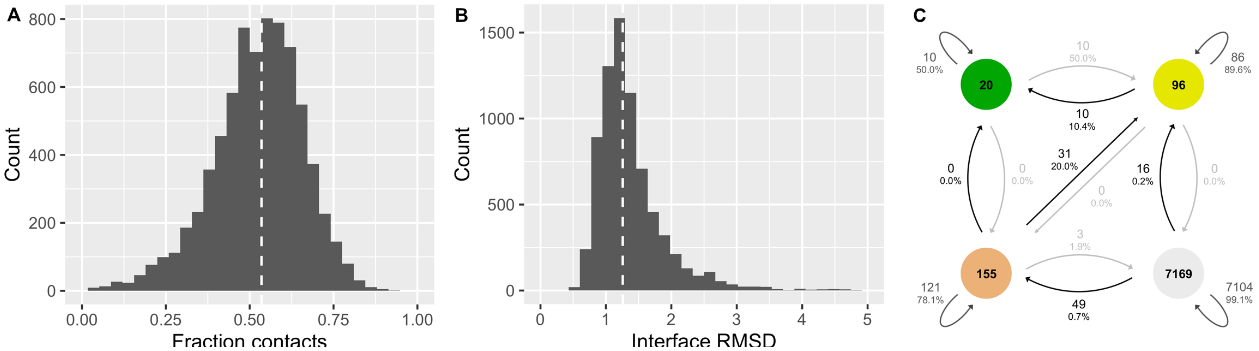


**Figure S1**. Quality assessment of the AlphaFold2-generated models in relation to its provided docking-generated template structure. Distribution in the fraction of conserved template contacts (**A**) and interface RMSD (**B**) between the AF2-generated model and its provided docking-generated template for the decoys in the unbound-backbone set. The RMSD calculations only include the Cα atoms. All decoys generated with ProPOSE, ZDOCK and PIPER were combined. The median of the distribution is indicated by the dashed white line. Transitions in model quality from the docking-generated model to the corresponding AF2-generated model (**C**). Transitions between the high-quality and incorrect classes were not observed. Colors denote structure quality levels as defined by CAPRI classification (Ref 35): high (green), medium (yellow), acceptable (beige) and incorrect (grey).

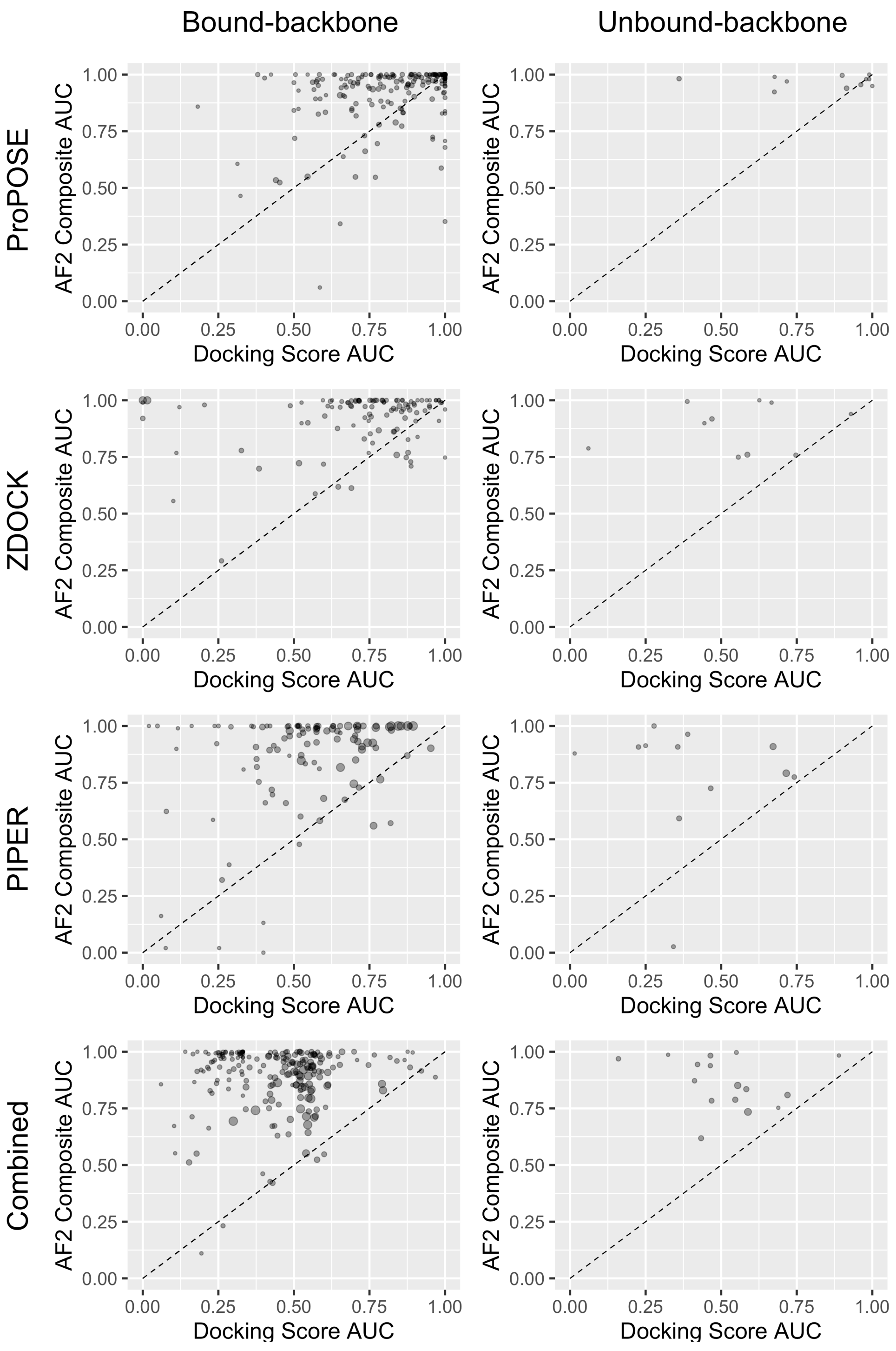


**Figure S2**. Classification of true positive and false positives according to AUC calculated from the docking scores compared to AF2-rescored for the bound-backbone and unbound-backbone sets. A medium-quality model is required for establishing success as opposed to an acceptable-quality (**Figure 2**). The AF2-generated models were used for success attribution. The number of data points in each plot are 199, 11, 109, 10, 109, 12, 211 and 16 (from left to right and top to bottom). AlphaFold2 improves the classification for 131 (66%), 7 (64%), 96 (88%), 10 (100%), 98 (90%), 11 (92%), 204 (97%) and 16 (100%).

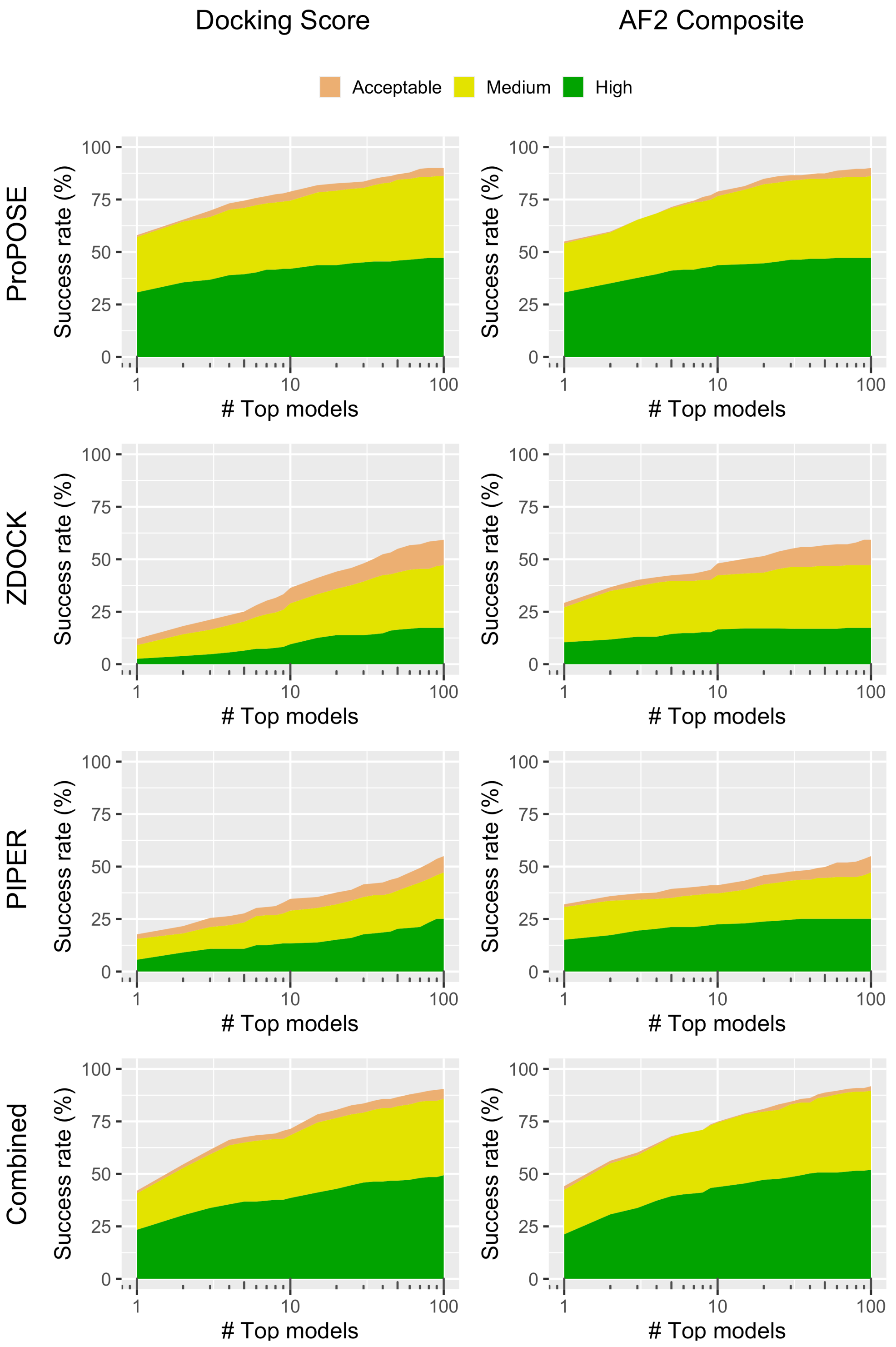


**Figure S3**. Success rates broken down for each model quality (Ref 35) for the docking methods as a function of the number of top models considered from the ranked-list. The rates are shown for models ranked using the docking scores and the $AF2_{Composite}$ score for the bound-backbone set. The rates were plotted on a logarithmic scale for better visibility. The success was evaluated using the AF2-generated models.

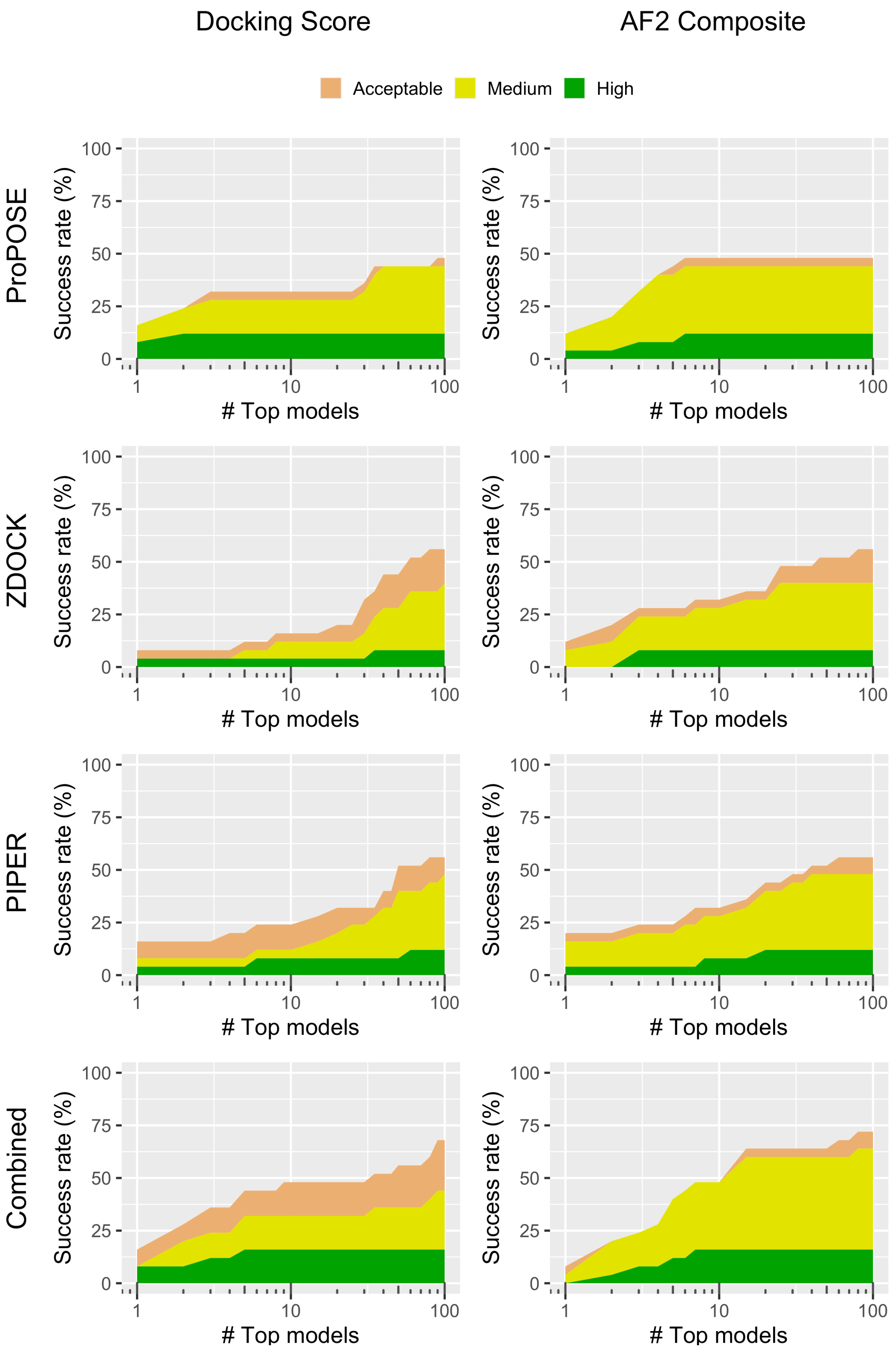


**Figure S4**. Success rates broken down for each model quality (Ref 35) for the docking methods as a function of the number of top models considered from the ranked-list. The rates are shown for models ranked using the docking scores and the $AF2_{Composite}$ score for the unbound-backbone set. The rates were plotted on a logarithmic scale for better visibility. The success was evaluated using the AF2-generated models.

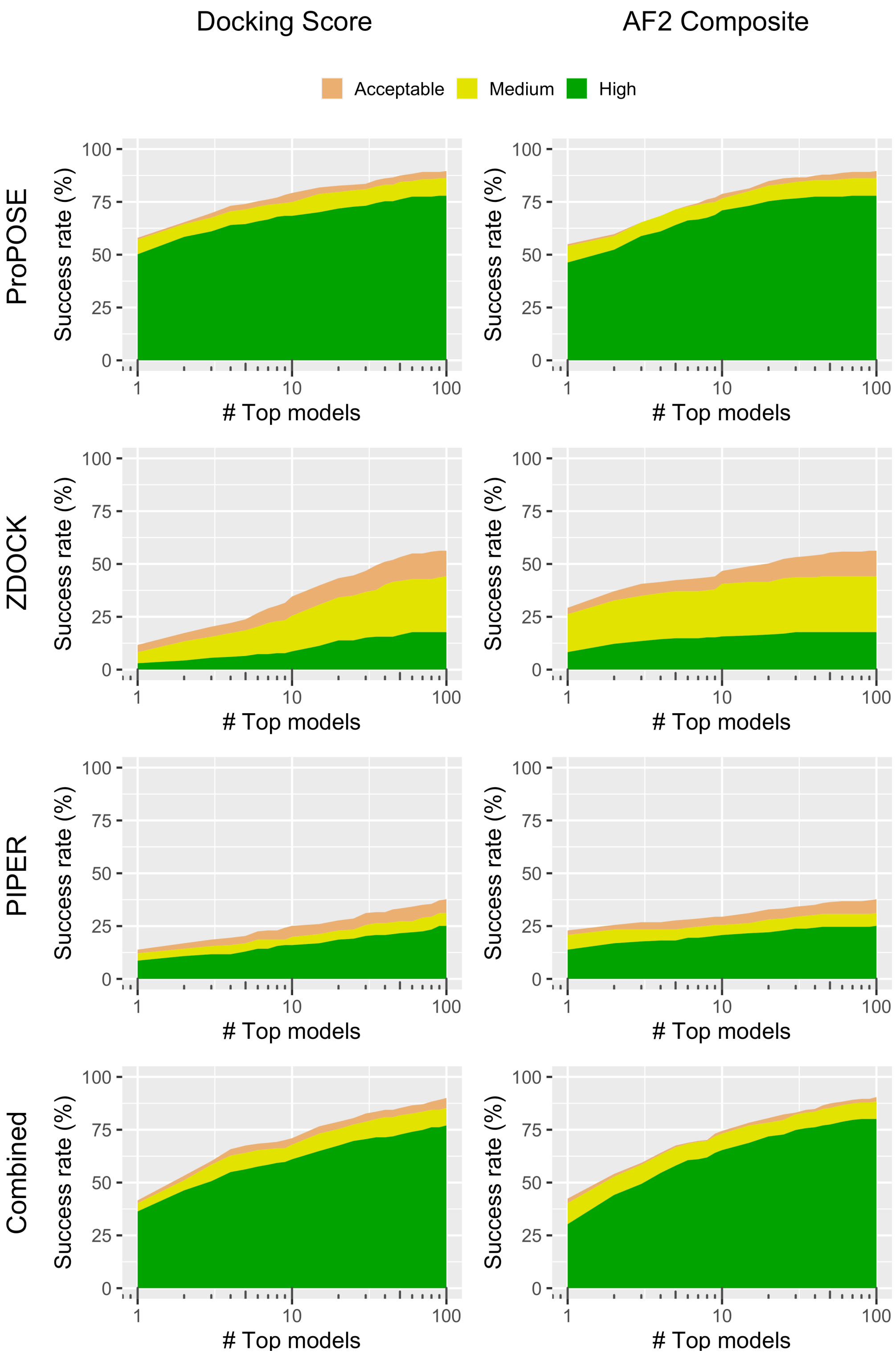


**Figure S5**. Success rates broken down for each model quality (Ref 35) for the docking methods as a function of the number of top models considered from the ranked-list. The rates are shown for models ranked using the docking scores and the $AF2_{Composite}$ score for the bound-backbone set. The rates were plotted on a logarithmic scale for better visibility. The success was evaluated using the docking-generated models.

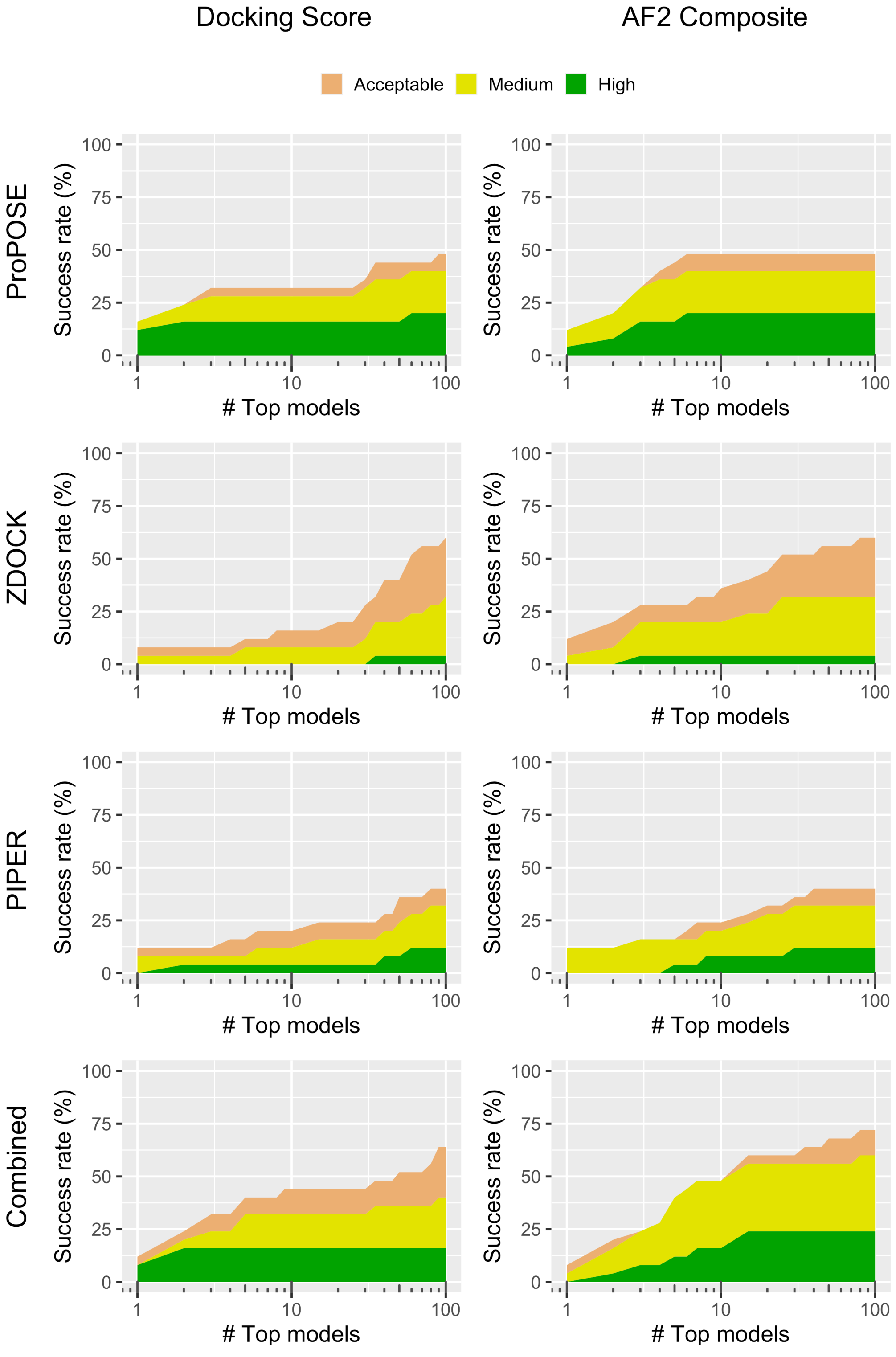


**Figure S6**. Success rates broken down for each model quality (Ref 35) for the docking methods as a function of the number of top models considered from the ranked-list. The rates are shown for models ranked using the docking scores and the AF2$_{Composite}$ score for the unbound-backbone set. The rates were plotted on a logarithmic scale for better visibility. The success was evaluated using the docking-generated models.

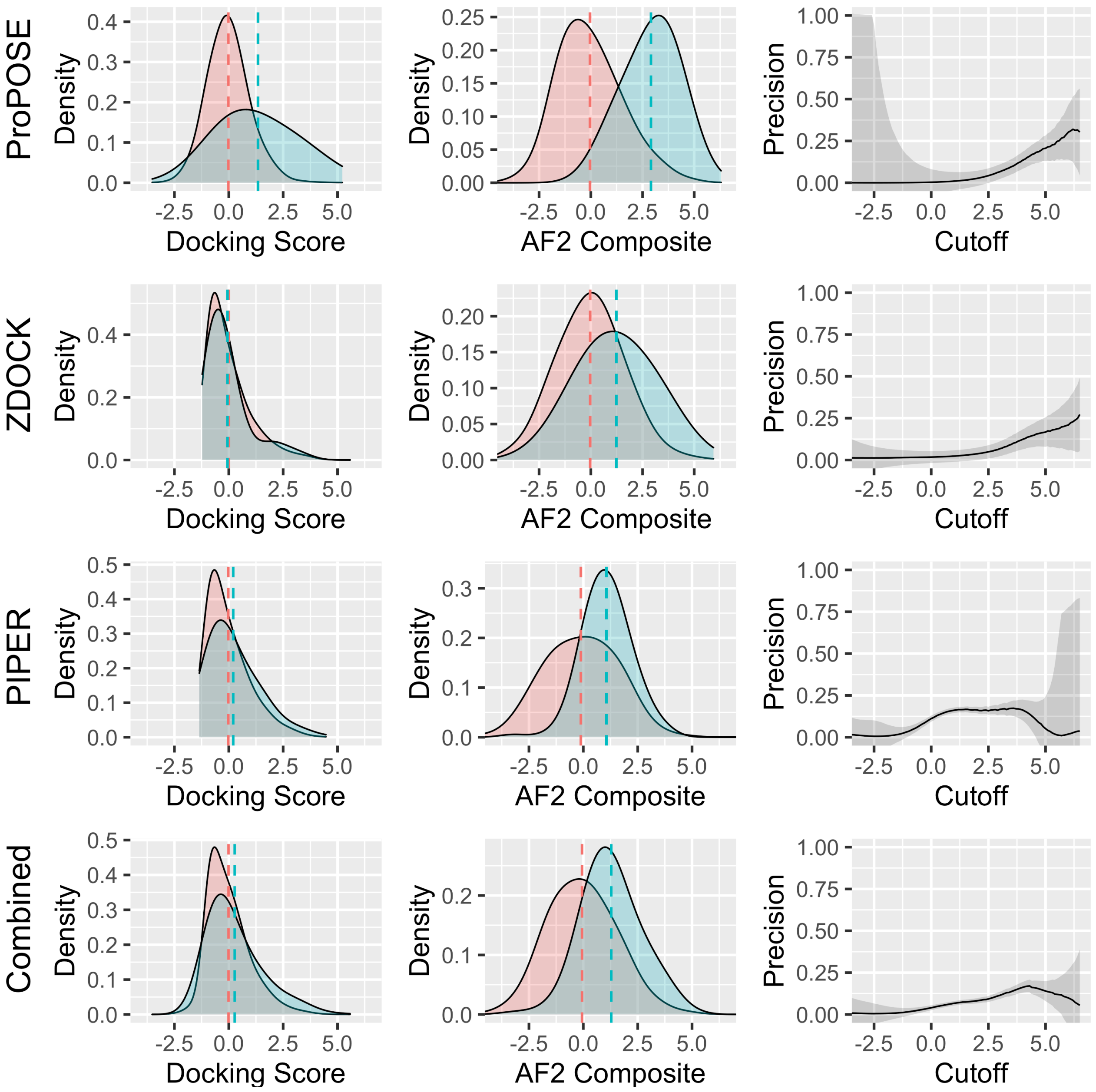


**Figure S7**. Density plots for the docking scores and the $AF2_{Composite}$ score for the negative and positive sets from the unbound-backbone set. The means of the distributions are marked with dashed lines. Precision curves were built from calculating the fraction in the number of true positive over the total number of true positives and false positives according to a given cutoff in the $AF2_{Composite}$ score. The smoothed densities were used to build the precision curves to avoid outlier bias. Shadings around the precision curves indicate the errors on these curves, which were estimated based on the number of data points, *i.e.* less data points incur larger errors.

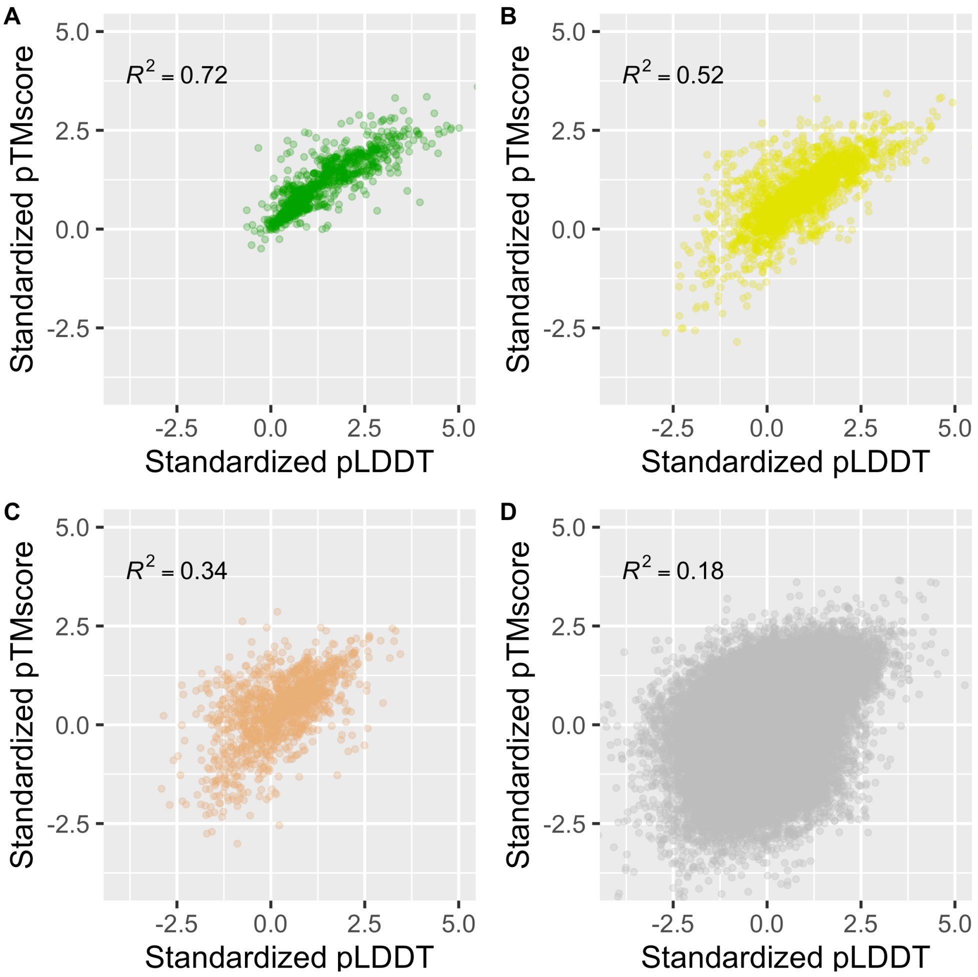


**Figure S8**. Correlation between the two components of the $AF2_{Composite}$ score. The scatterplots are shown separately for the four CAPRI model quality levels (Ref 35): high (**A**), medium (**B**), acceptable (**C**) and incorrect (**D**). All models generated by ProPOSE, ZDOCK and PIPER on the bound-backbone set are shown.

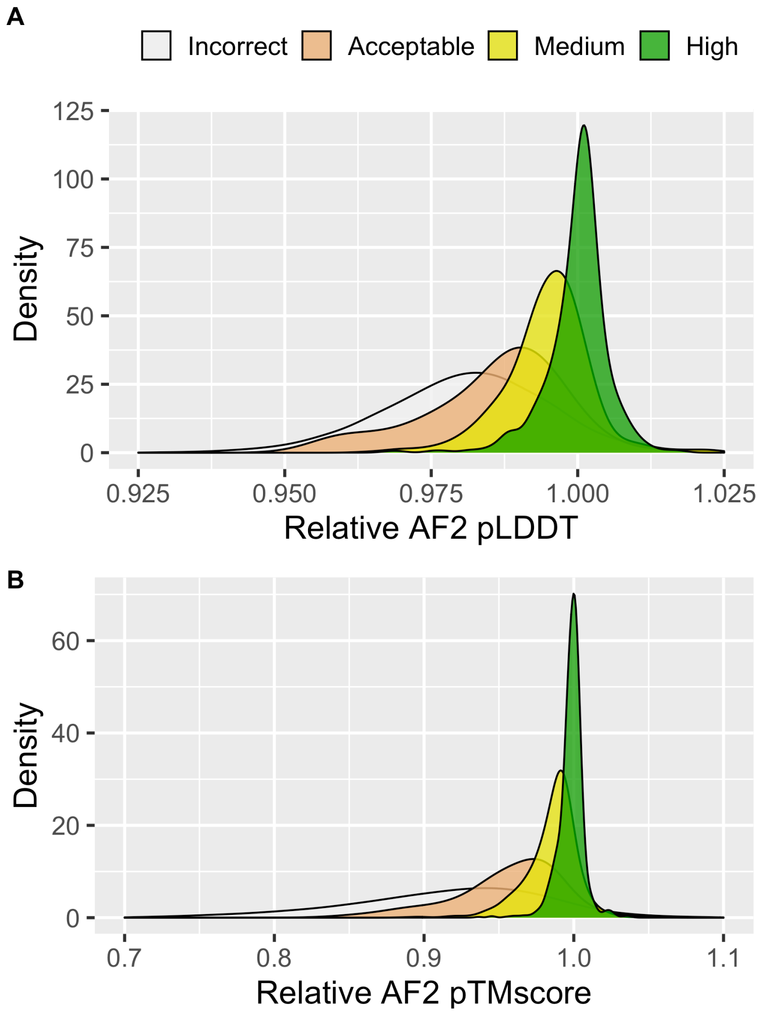


**Figure S9**. Smoothed density distribution of the relative AF2 confidence scores for the bound-backbone models in the negative (incorrect) and positive (acceptable-, medium- and high-quality) sets. The confidence scores are relative to those obtained by AlphaFold2 when provided with the crystal structures as input. The absolute pTMscore (**A**) and pLDDT (**B**) values are used as reference confidence scores for comparison due to the inability of deriving the composite score from a single structure in the case of the crystal. The confidence scores decrease as the quality of the model degrades.

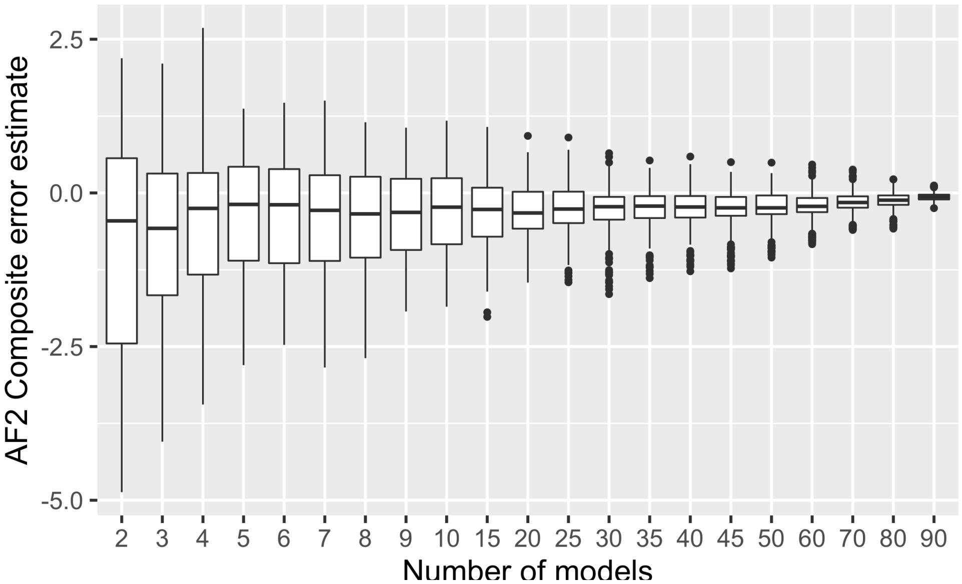


**Figure S10**. Error estimate on the AF2$_{\text{Composite}}$ as a function of the number of docking-generated models per system, *i.e.* the ensemble size. The error is calculated by subtracting the AF2$_{\text{Composite}}$ scores from models collected from smaller samples (ensembles of size below 100) to the theoretical AF2$_{\text{Composite}}$ scores from the population (ensemble of size 100). The unbound-backbone ProPOSE-generated models set were used for plotting. As the size of the ensemble increases, the estimate values approach those of the population.
